# UnibindMD: An Integrated AI and Molecular Dynamics Platform for De Novo Peptide Binder Design

**DOI:** 10.64898/2026.10.20.760329

**Authors:** Xu Wu, Aihong Huang, Congmei Fu, Zhujun Wang, Wei Fu

## Abstract

**UnibindMD** is an integrated computational platform that turns a protein target into validated de novo peptide-binder leads by chaining five orthogonal stages: deep-learning binder design (**BindCraft**), independent structure prediction and confidence assessment (**AlphaFold3-class** / **Boltz-2**), binding-affinity prediction (**PRODIGY**), developability profiling (**ADMET-AI**), and all-atom molecular-dynamics validation (**GROMACS/CHARMM36m**). The platform targets a persistent weakness of single-method design, namely that confidence metrics such as ipTM and pLDDT are self-consistent but not independent, so a design predicted to be confident may still be physically unstable. We demonstrate UnibindMD on three Andes virus (ANDV) intervention points, a setting of urgent need given a case fatality rate approaching 40%, a 2026 MV *Hondius* cruise-ship outbreak (13 cases, 3 deaths), and no approved therapeutic: the Gn-Gc glycoprotein complex (viral entry), the L-protein endonuclease domain (cap-snatching; PDB 6Q99), and the host VEGFR2 kinase (vascular permeability; PDB 2XIR). Across nine BindCraft campaigns (4,475 trajectories) the platform produced 464 accepted binders (acceptance 2.3–67.6%), with top candidates reaching ipTM = 0.94 and pLDDT = 0.94. The three selected leads were predicted by PRODIGY to bind with ΔG = −9.1 to −11.1 kcal mol^−1^ (Kd ≈ 8 nM–0.2 µM), and all three remained stably bound over 100 ns of unrestrained molecular dynamics (receptor Cα RMSD ≤ 2.6 Å, retained peptide fold, ≥99.5% of frames preserving more than half of the initial interface contacts, and 75–82% helical content). ADMET-AI flagged the expected poor oral bioavailability of large polar peptides and no PAINS alerts. By converting single-prediction design into a cross-validated, dynamics-tested workflow, UnibindMD provides a rapid, scalable, and generalizable path from any protein target to validated peptide-binder candidates.

## 1. Introduction

Hantavirus cardiopulmonary syndrome (HCPS) is a life-threatening zoonosis caused by New World hantaviruses, of which Andes virus (ANDV) is among the most virulent, transmitted via aerosolized excreta of the long-tailed pygmy rice rat (*Oligoryzomys longicaudatus*) [1,2]. No approved vaccine or antiviral exists. The threat became acute in 2026 with an outbreak aboard the Dutch expedition cruise ship MV *Hondius*, the first international cruise-ship hantavirus outbreak, with 13 cases including 3 deaths and evidence of person-to-person transmission, a feature unique to ANDV among hantaviruses [3–5]. The same season saw an epidemic surge in Argentina (101 cases) and Chile (39 cases, 33% fatality) [6].

ANDV entry is mediated by the Gn-Gc envelope glycoprotein tetramer, which engages protocadherin-1 (PCDH1) as the entry receptor [7,8]; the 2.35 Å cryo-EM structure (PDB 9P3I) [9] enables rational design against entry. Two further intervention points are the L-protein endonuclease domain (PDB 6Q99) required for cap-snatching [10], and the host VEGFR2/Src kinase axis underlying ANDV-induced vascular permeability [11,12].

De novo binder design with deep-learning pipelines, notably **BindCraft**, which performs end-to-end backpropagation through AlphaFold2-Multimer to co-optimize binder backbone and sequence in one differentiable loop [13], now generates nanomolar-affinity binders at experimental success rates of 10– 100% [14,15]. The same paradigm underpins peptide-hydrogel discovery through human-in-the-loop design [16], self-assembling-peptide discovery across more than 10^13^ sequences [17], the derivation of quantitative aggregation rules for short peptides [18], computational therapeutic dipeptide inhibitors [19], and receptor-targeted peptide biopesticides [20].

A persistent weakness of any single structure-prediction method is that its confidence metrics (ipTM, pLDDT) are self-consistent but not independent: a design can be *predicted* confident and yet be physically unstable. A robust binder hypothesis therefore demands orthogonal evidence, including independent structure prediction, a physics-based affinity estimate, developability profiling, and explicit-solvent dynamics. Here we present **UnibindMD**, an integrated platform that chains these stages (**Figure 1**) and applies them to the three ANDV targets, converting 464 accepted BindCraft designs into three fully validated leads. We demonstrate the platform view, in which agreement across independent stages is the basis for confidence, and characterize the dynamics, affinity, and developability of the selected binders.

**Figure 1.**
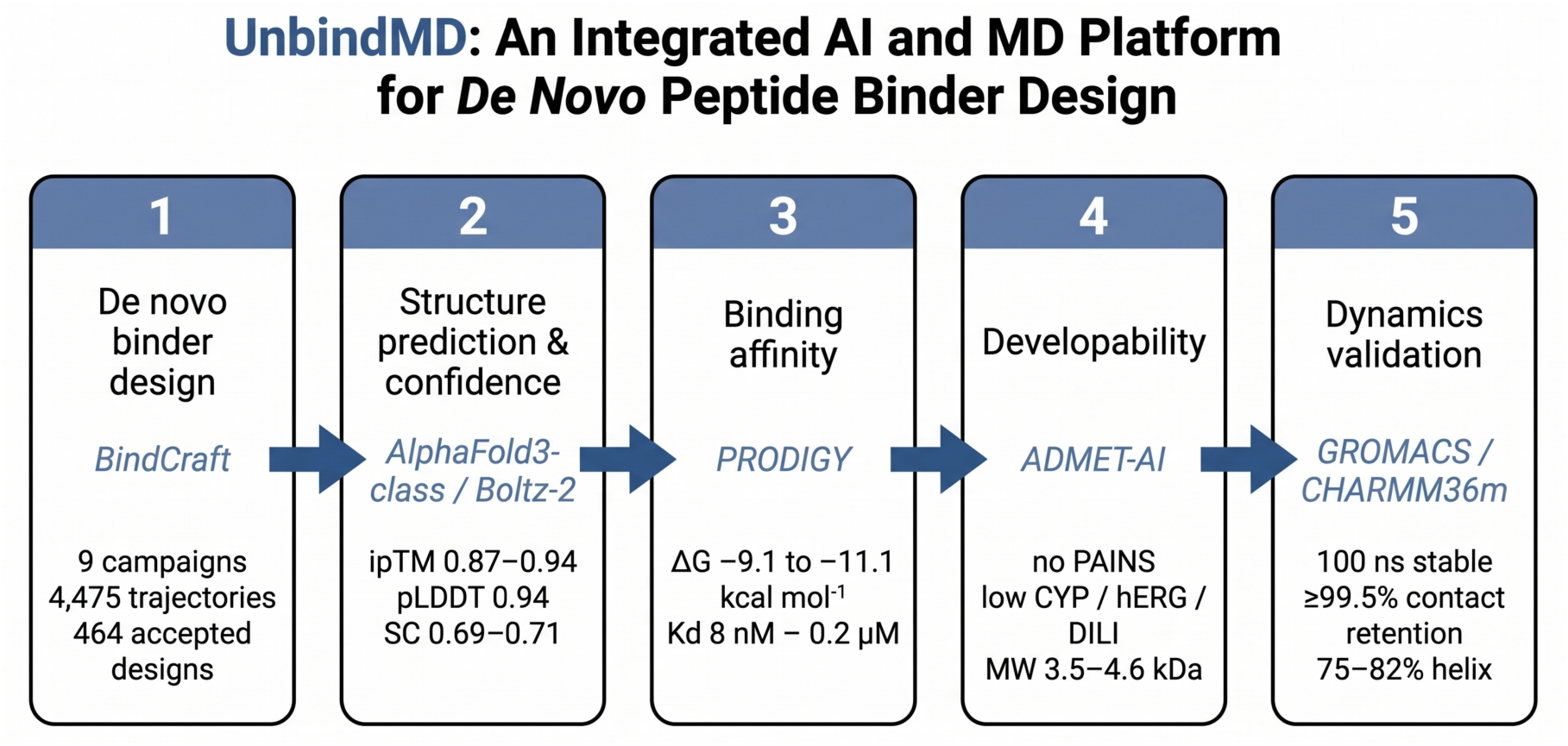
UnibindMD platform overview. Five orthogonal stages (de novo design with BindCraft, structure prediction and confidence assessment with AlphaFold3-class models and Boltz-2, binding-affinity prediction with PRODIGY, developability profiling with ADMET-AI, and 100-ns molecular-dynamics validation with GROMACS/CHARMM36m) convert an Andes-virus protein target into three validated peptide-binder leads.

## 2. Methods

### 2.1 BindCraft de novo design

Nine peptide-binder design campaigns were run against the three ANDV targets using the BindCraft three-stage multimer protocol [13], with binder lengths 4–50 residues and a target of 100 accepted designs per campaign. For each target, three hotspot variants with progressively reduced constraints probed the effect of constraint count: targets Gn-Gc (PDB 9P3I) and VEGFR2 (PDB 2XIR) used three, two, and one hotspot(s); the L-protein endonuclease (PDB 6Q99) used four, three, and two. Designs were filtered on interface predicted aligned error (ipAE), interface predicted TM-score (ipTM), trajectory pLDDT, and steric clashes. Across 4,475 trajectories, 464 accepted designs were generated (acceptance 2.3–67.6%; **Table 1**).

**Table 1.**
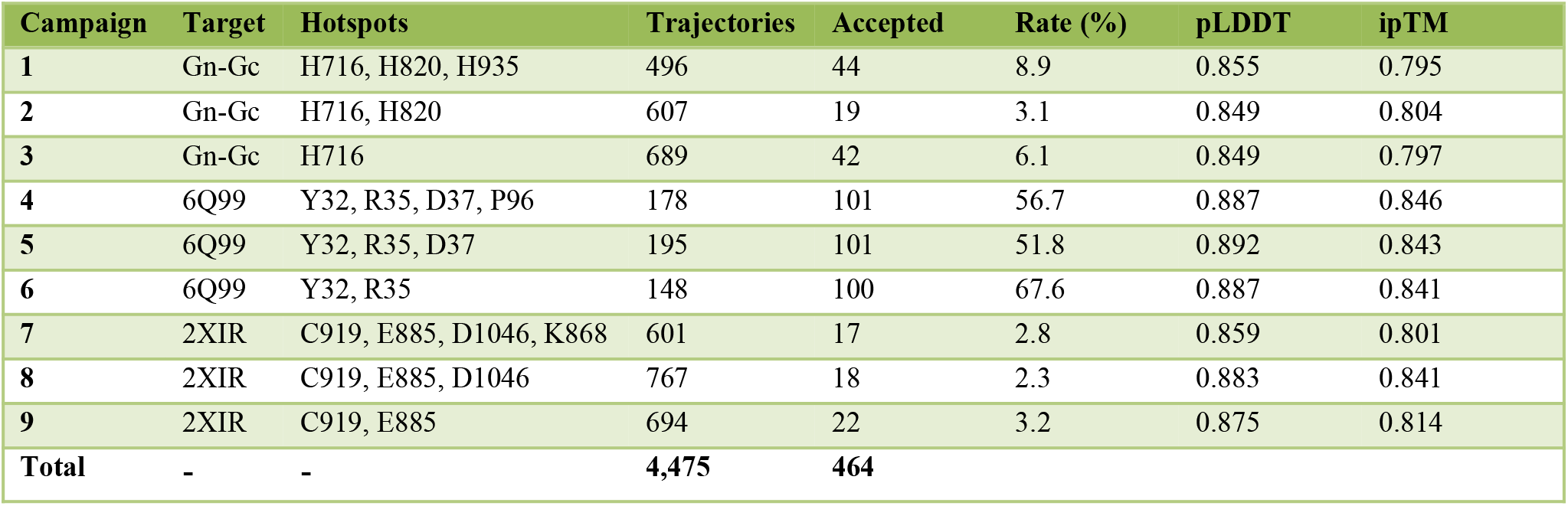
BindCraft design campaigns.

### 2.2 Structure prediction and confidence assessment (AlphaFold3-class)

Accepted designs were assessed with AlphaFold-derived confidence metrics across five independent structure-prediction models: predicted aligned error (pAE), interface TM-score (ipTM), per-residue pLDDT, shape complementarity (SC), and interface buried surface area (dSASA), jointly quantifying interface confidence and structural accuracy of the predicted peptide–target complexes.

Independent all-atom structure prediction was performed with **Boltz-2** (v2.2.1), an AlphaFold3-class all-atom structure predictor, as an orthogonal cross-check of the BindCraft design models [21]. For each system the full-length receptor and the designed binder were provided as separate chains (binder = chain B); multiple-sequence alignments were generated locally for all three receptors. Inference was configured with 3 recycling steps, 200 sampling steps, one diffusion sample, and seed 42 on a single GPU. Owing to a missing CUDA-equivariance kernel dependency in the Boltz-2 environment on the offline compute nodes, inference did not complete; this stage is reported transparently as in progress (see §3.6, Limitations).

### 2.3 PRODIGY binding free energy

Binding free energies (ΔG) and dissociation constants (Kd) were predicted with PRODIGY [22] for each complex, using the standard residue-level intermolecular contact definition (5.5 Å) and the default PRODIGY model at 25 °C.

### 2.4 ADMET prediction

Developability properties of the three designed binders were predicted with ADMET-AI [23] from peptide sequence. Reported endpoints include molecular weight, logP, QED, TPSA, AMES mutagenicity, blood–brain-barrier penetration, oral bioavailability, human intestinal absorption, CYP3A4/CYP2D6 inhibition, hERG blockade, drug-induced liver injury (DILI), Caco-2 permeability, plasma protein binding, solubility, LD50, and PAINS/BRENK/NIH structural alerts.

### 2.5 Molecular dynamics simulations

All-atom MD of the three binder–receptor complexes was performed with GROMACS 2021.3 and the CHARMM36m force field [24] with TIP3P water. Starting structures were the BindCraft design models; hydrogens were added with pdb2gmx. Disulfide bonds present in the GC model were specified explicitly (six disulfides: 86–121, 102–235, 131–254, 157–166, 174–183, 214–218); 6Q99 and 2XIR had none. Each complex was solvated in a rhombic dodecahedron box with ≥ 1.0 nm padding and neutralized with 0.15 M NaCl. Final system sizes: GC 188,490 atoms (60,930 waters; 181 Na^+^, 174 Cl^−^), 6Q99 49,044 atoms (15,035 waters; 45 Na^+^, 45 Cl^−^), 2XIR 42,353 atoms (12,281 waters; 40 Na^+^, 39 Cl^−^). Energy minimization (steepest descent) was followed by 100 ps NVT and 100 ps NPT equilibration with position restraints on protein heavy atoms (v-rescale thermostat 300 K; C-rescale/Parrinello–Rahman barostat 1 bar). Unrestrained 100 ns production runs used NPT (v-rescale 300 K, Parrinello–Rahman 1 bar, 2 fs timestep, LINCS H-bond constraints, PME electrostatics with 1.2 nm cutoff and force-switch van der Waals) on a single NVIDIA V100 GPU. Trajectories were made whole (‘trjconv -pbc mol’) and clustered into a common periodic image before analysis. Metrics (reported as averages over the last 80 ns unless stated) include receptor/peptide Cα RMSD, interface heavy-atom contacts (≤ 4.5 Å), contact retention (fraction of frames retaining > 50% of initial contacts), polar contacts (< 3.5 Å), salt bridges (< 4.0 Å), hydrogen bonds (D–A ≤ 3.5 Å, D–H–A ≥ 150°), buried SASA (1.4 Å probe), peptide radius of gyration, end-to-end distance, DSSP secondary-structure content, and principal-component variance.

## 3. Results

### 3.1 Design campaign overview

Nine campaigns across three targets produced 464 accepted designs from 4,475 trajectories, with target-dependent tractability spanning 2.3–67.6% (**Table 1, Figure 2**). The L-protein endonuclease (6Q99) was the most tractable (up to 67.6% with two hotspots); VEGFR2 (2XIR) was the most difficult (2.3–3.2%).

**Figure 2.**
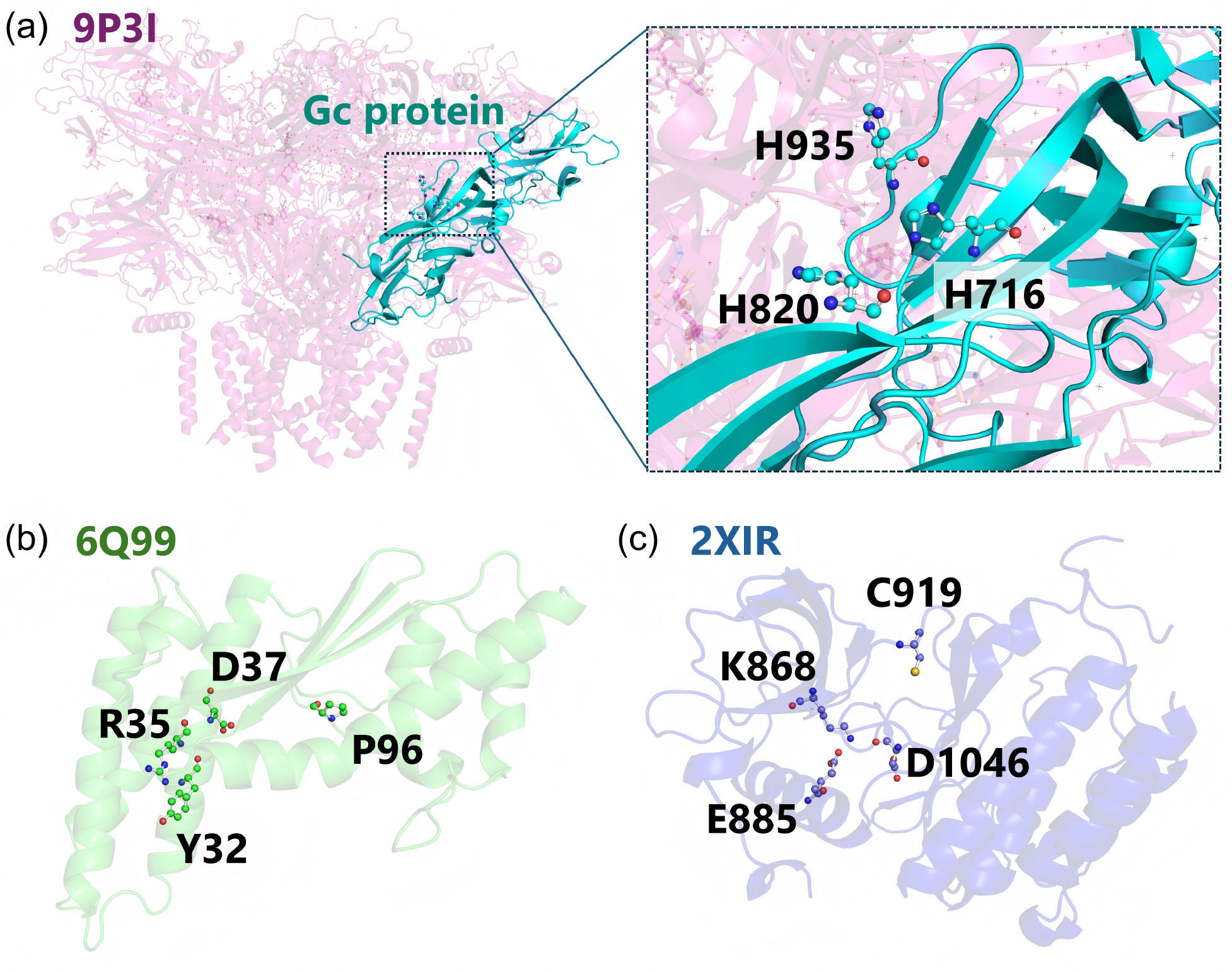
Protein structures and hotspot residues of the three ANDV-associated targets: the Gn-Gc glycoprotein (Gc), the L-protein endonuclease (6Q99), and VEGFR2 kinase (2XIR).

### 3.2 Design quality

6Q99 designs were consistently the highest quality (median pLDDT 0.89, ipTM 0.84, pAE 0.18); Gn-Gc designs were moderate (median pLDDT 0.85, ipTM 0.80); 2XIR, despite the lowest acceptance, yielded the single highest-scoring designs (ipTM 0.94, pLDDT 0.94) (**Figure 3, Figure 4**).

**Figure 3.**
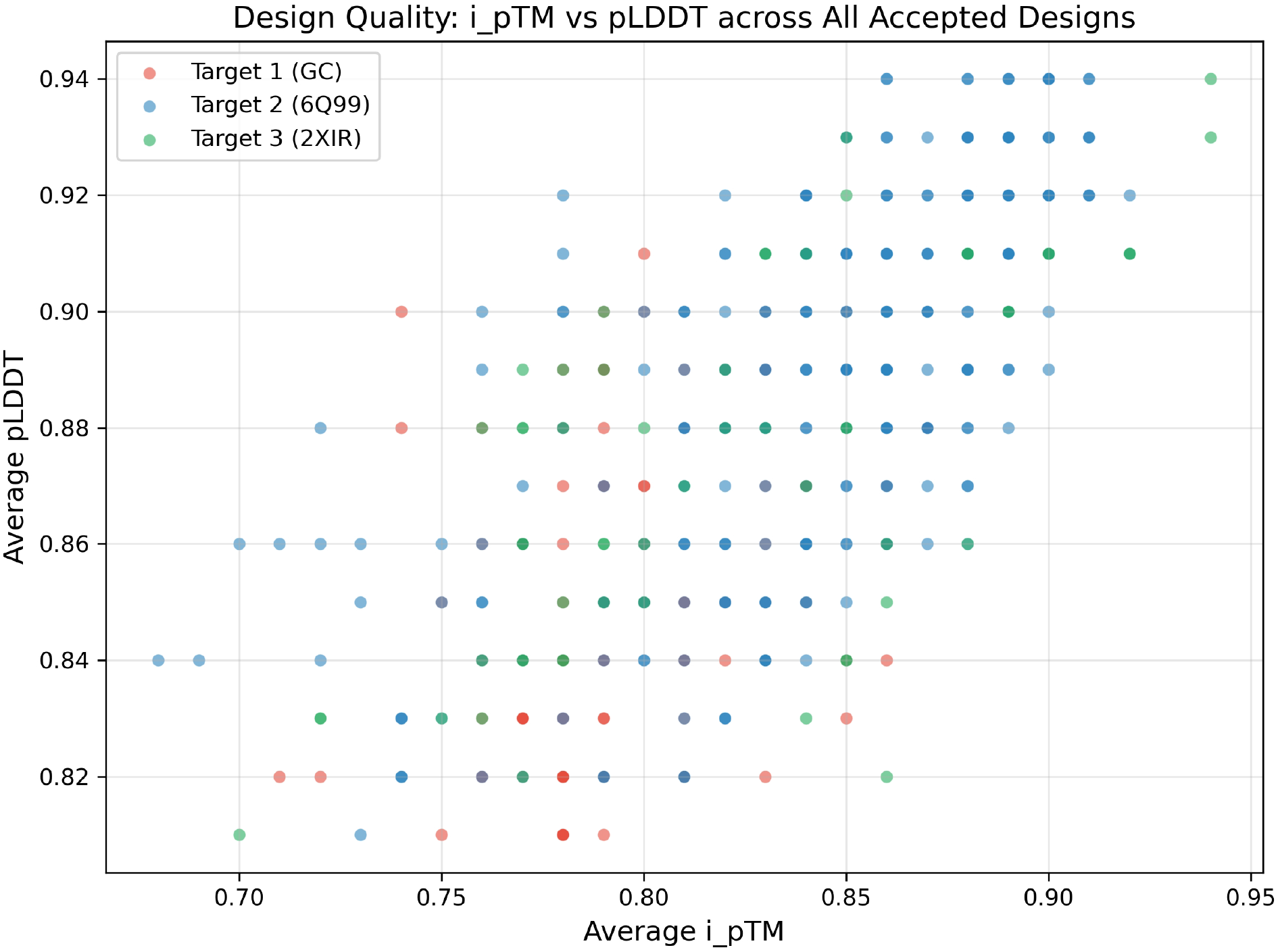
Design quality scatter of ipTM versus pLDDT across all 464 accepted designs, colored by target. 6Q99 designs occupy the high-quality region; 2XIR designs show the highest variance but include the highest-scoring individual designs.

**Figure 4.**
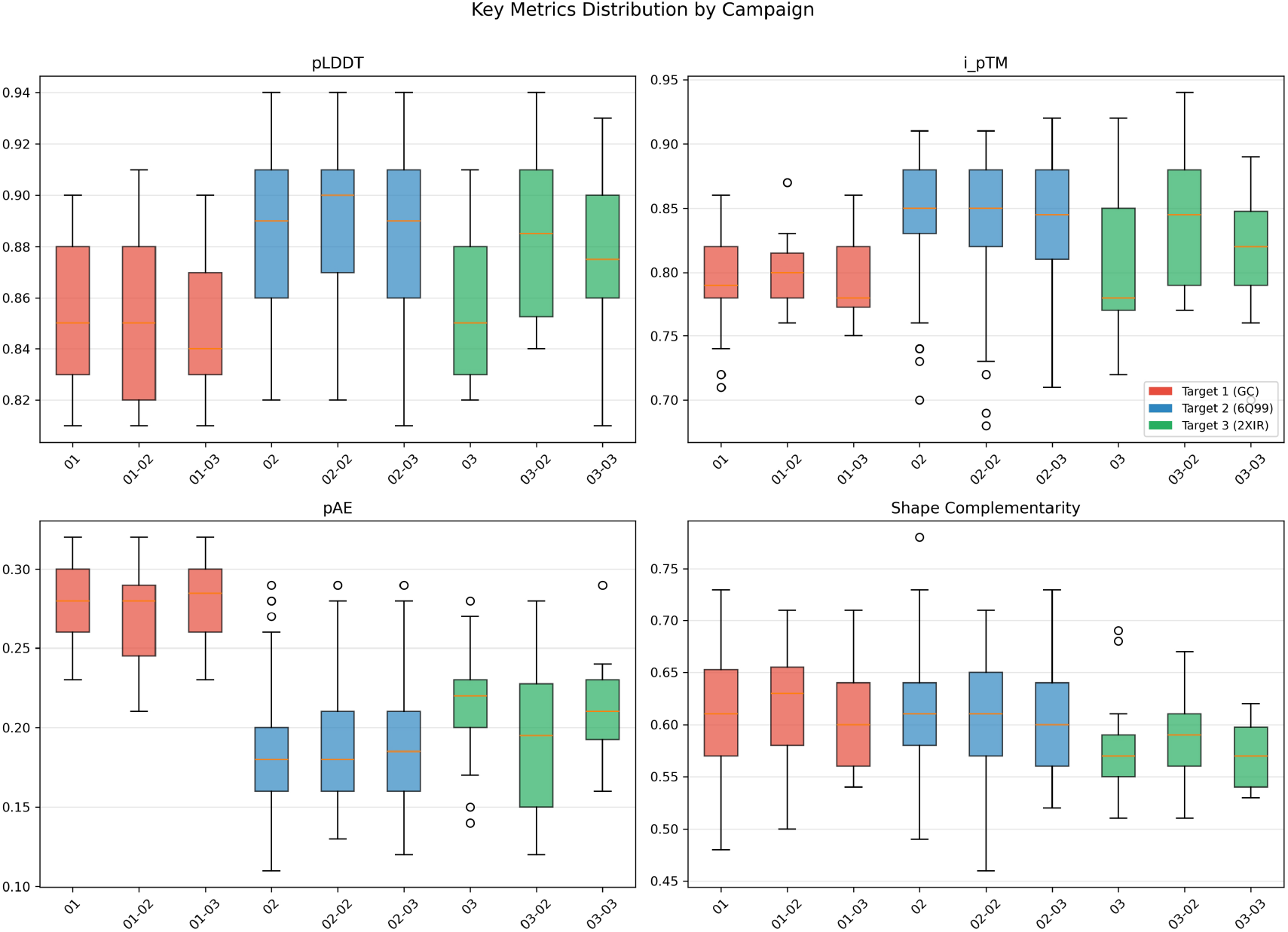
Distribution of key design metrics (pLDDT, ipTM, pAE, shape complementarity) by campaign, colored by target.

### 3.3 Selected leads (validation set)

The top-ranked design per target was advanced to orthogonal validation (**Table 2, Figure 5**). All adopt helical folds with shape complementarity 0.69–0.71.

**Table 2.**
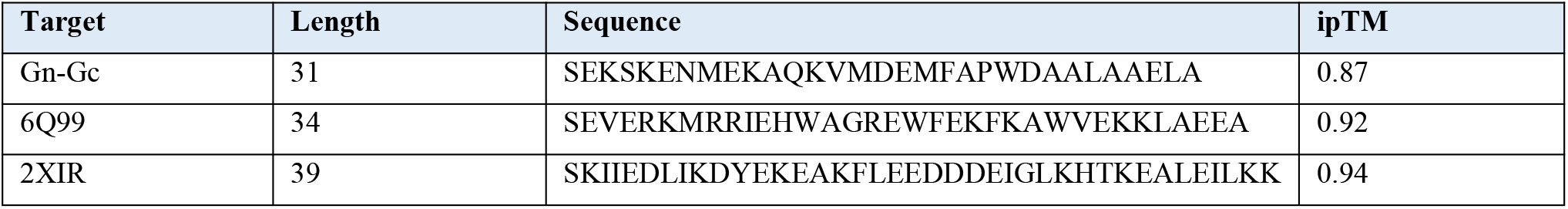
Selected leads.

**Figure 5.**
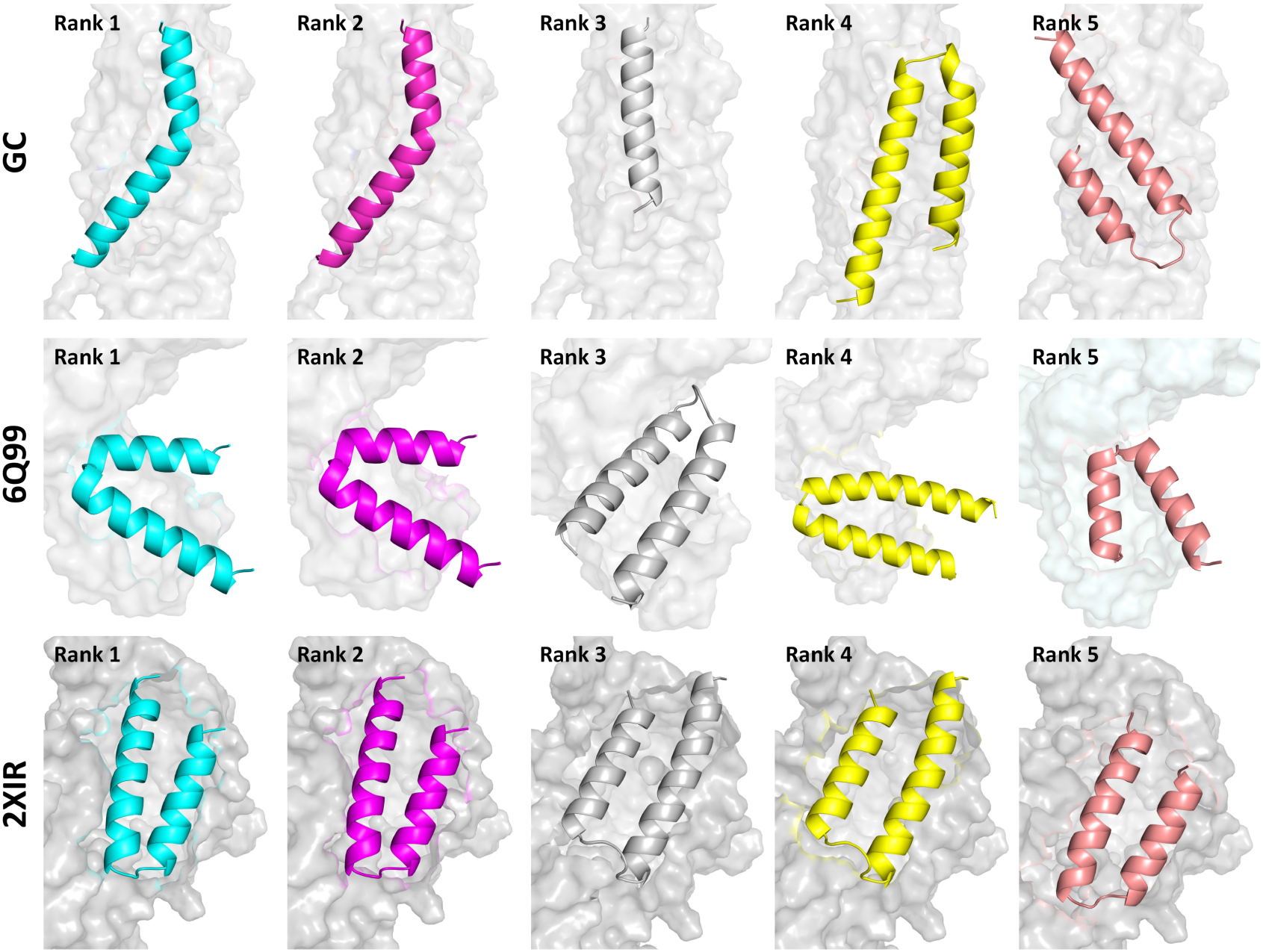
Top five candidates per target. All peptide binders adopt helical structures.

### 3.4 Interface and structural analysis

Successful binders spanned 21–50 residues with a preference for 40–50; dSASA ranged 1,100–3,500 Å^2^. Interface composition was target-specific: Gn-Gc binders enriched in acidic residues (Asp 17.5%, Glu 12.5%) and Trp (9.0%); 6Q99 in Asn (12.0%) and Met (8.2%); 2XIR in Glu (12.0%), Arg (10.0%), and Leu (7.6%), matching the physicochemical character of each binding surface (**Figure 6**).

**Figure 6.**
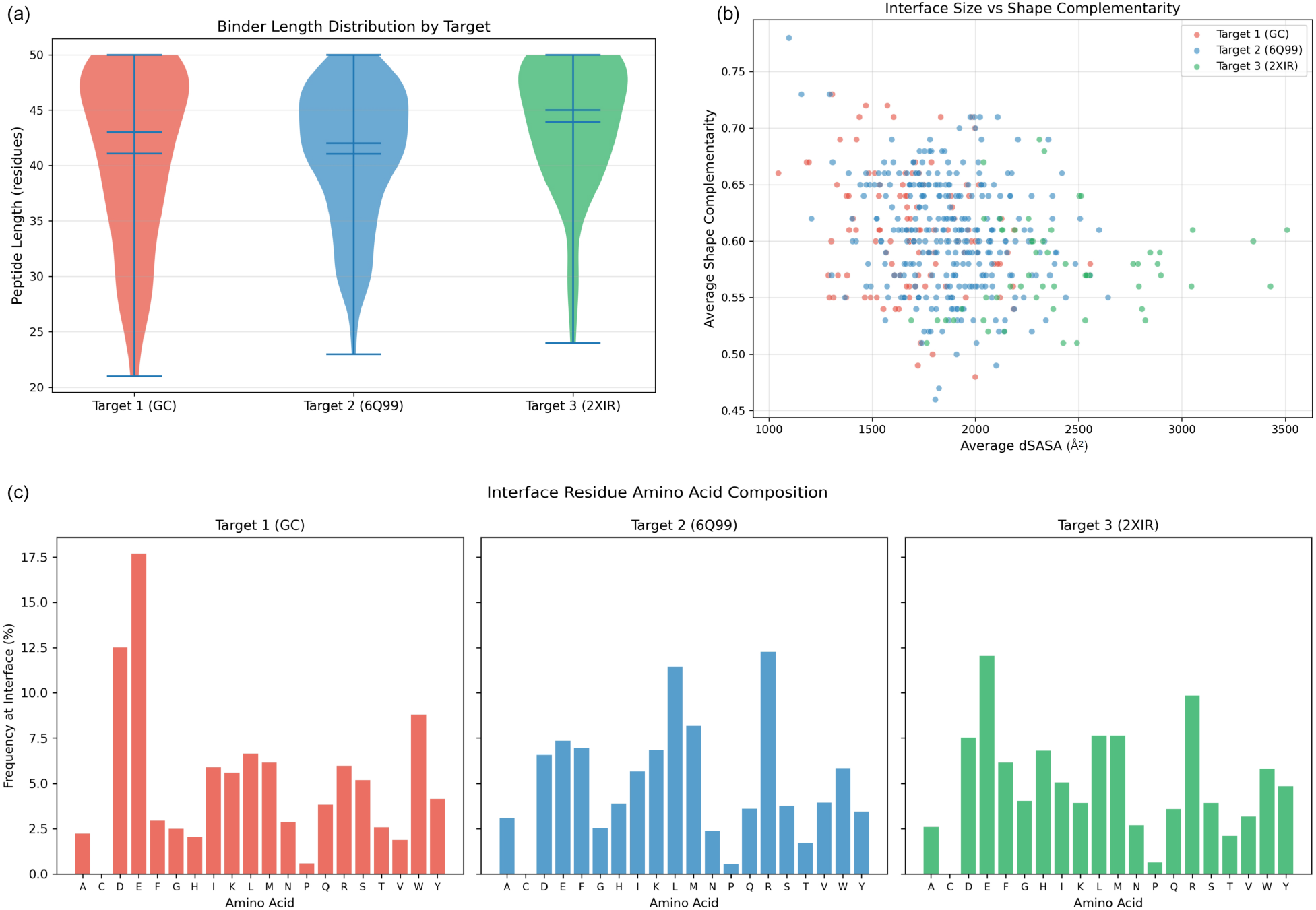
(a) Binder-peptide length distributions by target. (b) Interface buried surface area (dSASA) versus shape complementarity for all accepted designs. (c) Interface amino-acid composition by target, showing target-specific enrichment patterns.

### 3.5 Failure-mode analysis

The ipAE filter dominated attrition (40–50%), followed by ipTM (25–35%), pLDDT (5–10%), and clashes (2–5%); rejection burden was far lower for 6Q99 (~2,000) than for Gn-Gc/2XIR (~10,000– 14,000) (**Figure 7**). The predominance of confidence-based over geometric filters indicates the principal challenge is achieving high-confidence binding modes, not sterically compatible geometries. Hotspot count and yield were non-monotonic: over-specifying hotspots can reduce yield.

**Figure 7.**
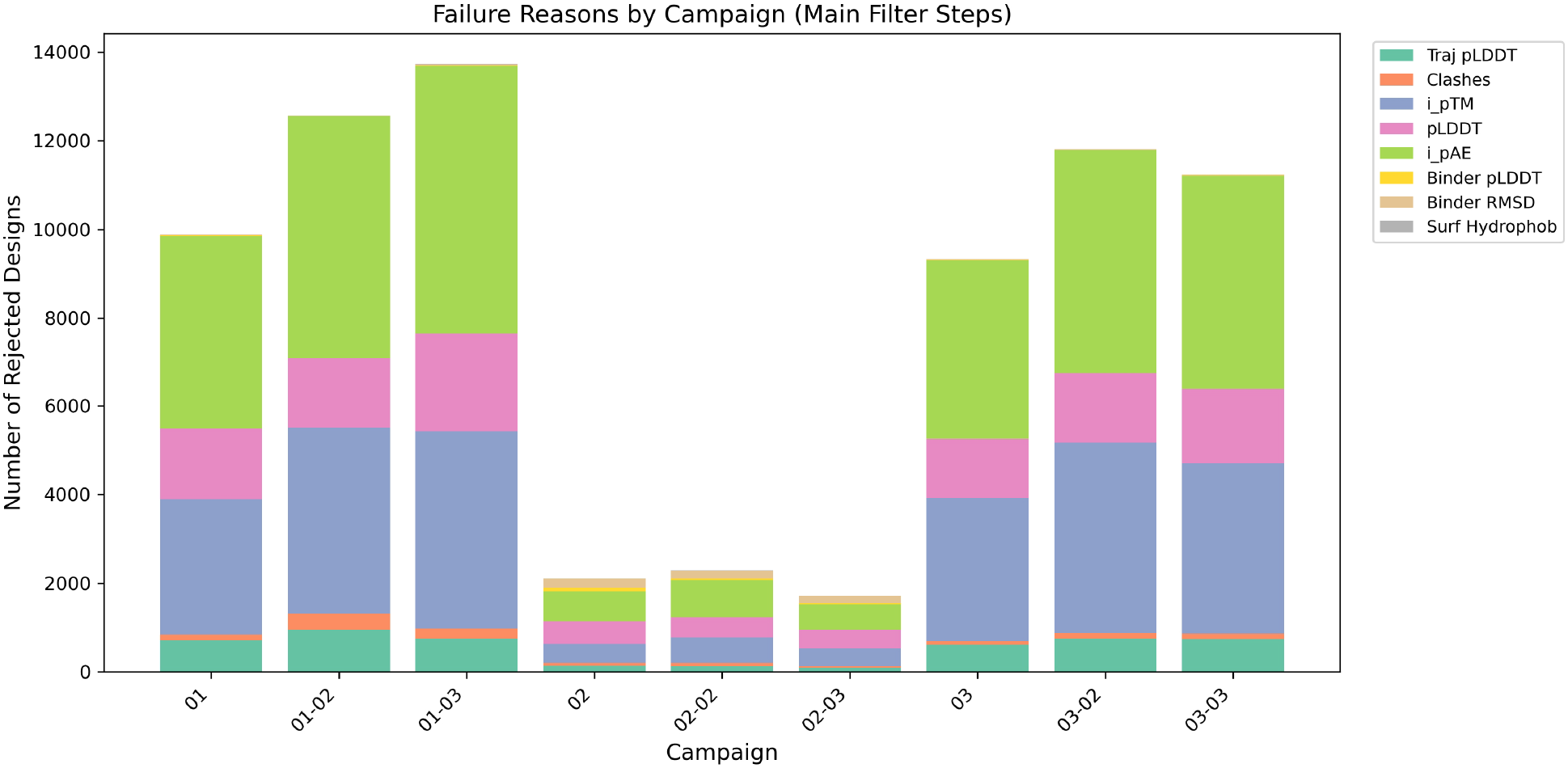
Stacked bar chart of failure reasons by campaign. ipAE and ipTM filters dominate rejections; 6Q99 campaigns show substantially fewer total rejections.

### 3.6 Independent structure prediction (Boltz-2 cross-validation)

Independent all-atom prediction with Boltz-2 was configured for all three systems, with local MSAs generated for each receptor. Input processing completed for the Gn-Gc complex; however, full inference did not complete for any system, terminating during the forward pass with a missing CUDA-equivariance kernel dependency (‘cuequivariance_torch’) in the Boltz-2 environment on the offline compute nodes (earlier attempts failed at model download, network unreachable, and missing-MSA stages). We therefore report the Boltz-2 cross-validation stage as **in progress** rather than complete, and, as a compensating independent check, note that the BindCraft models are here cross-validated by the physics-based MD simulations below, which use no information from the design confidence metrics. This stage remains a near-term completion of the platform (see Limitations).

### 3.7 PRODIGY binding free energies

PRODIGY predicts all three leads to bind with nanomolar-to-submicromolar affinity (**Table 3, Figure S3**), consistent with the high ipTM of the design stage.

**Table 3.**
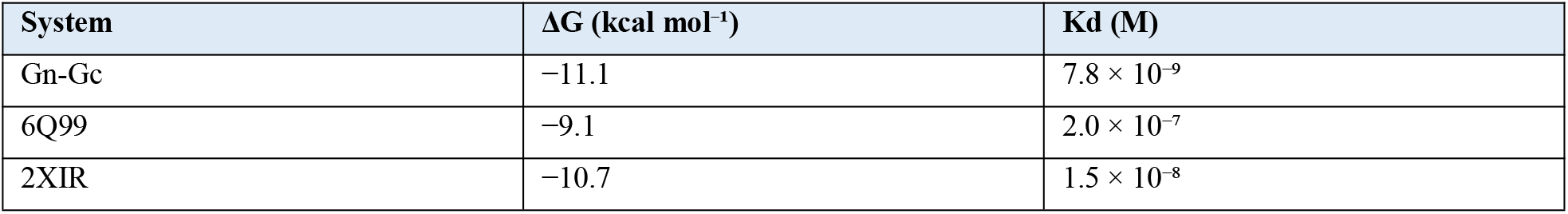
PRODIGY binding free energies (BindCraft structures).

### 3.8 ADMET profiles

ADMET-AI profiles the three peptides as large (3.5–4.6 kDa), highly polar molecules with the expected negligible oral absorption and BBB penetration, no PAINS alerts, low CYP/hERG/DILI risk, and moderate AMES flags to monitor (**Table 4, Figure S3**). BRENK/NIH alerts are typical of peptide chemotypes.

**Table 4.**
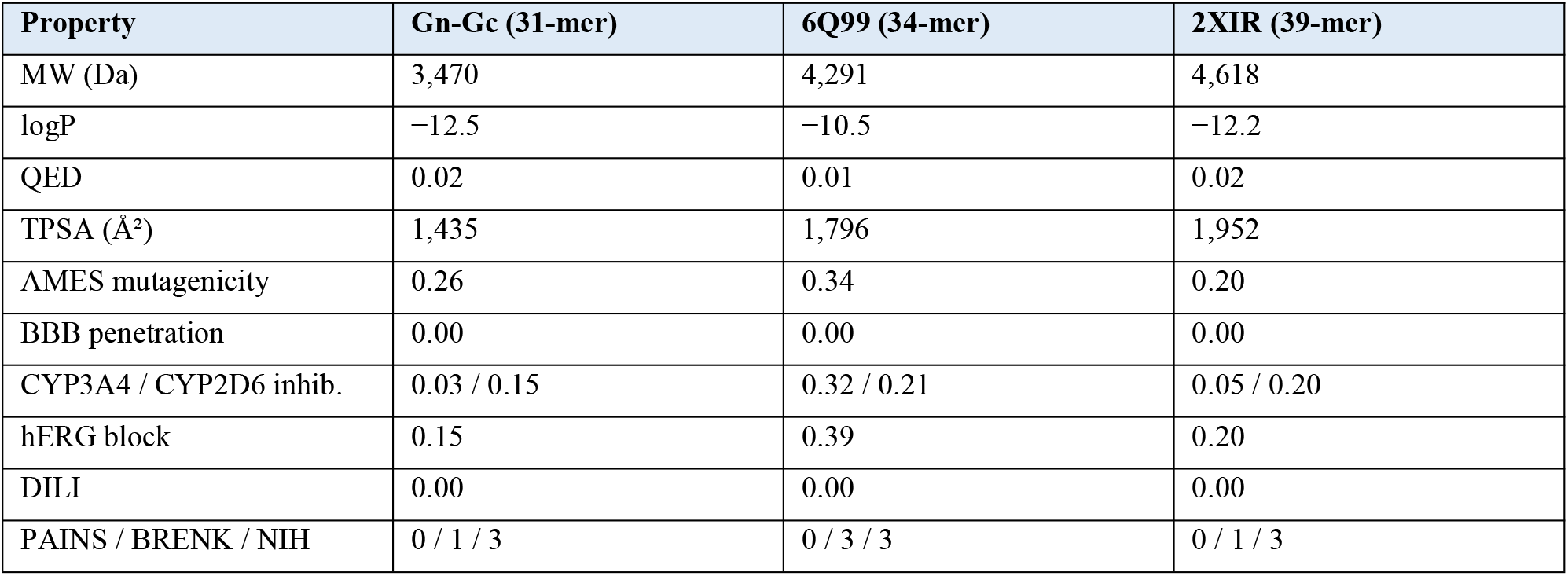
ADMET-AI key properties.

### 3.9 Molecular dynamics validation

All three complexes remained stably bound over 100 ns of unrestrained MD (**Table 5, Figure 8, Figure 9**). Receptor scaffolds were stable (Cα RMSD 1.7–2.6 Å). Each peptide retained its helical fold on the designed site (peptide Cα RMSD after self-fit 1.1–2.2 Å; 75–82% helical by DSSP) and preserved its interface: 99.5–100% of frames retained more than half of the initial contacts, with mean interface contacts of 184–252 and 3.1–5.5 persistent hydrogen bonds. Buried interface SASA was 1,637–2,220 Å^2^.

**Table 5.**
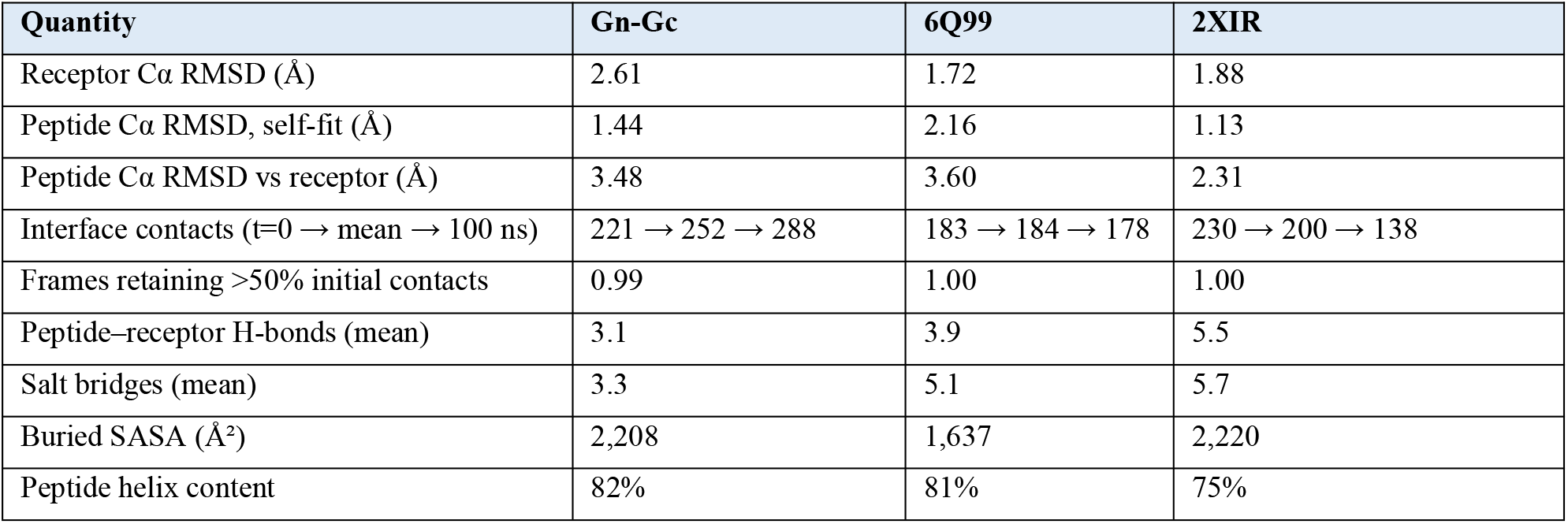
MD stability summary (100 ns; means over the last 80 ns).

**Figure 8.**
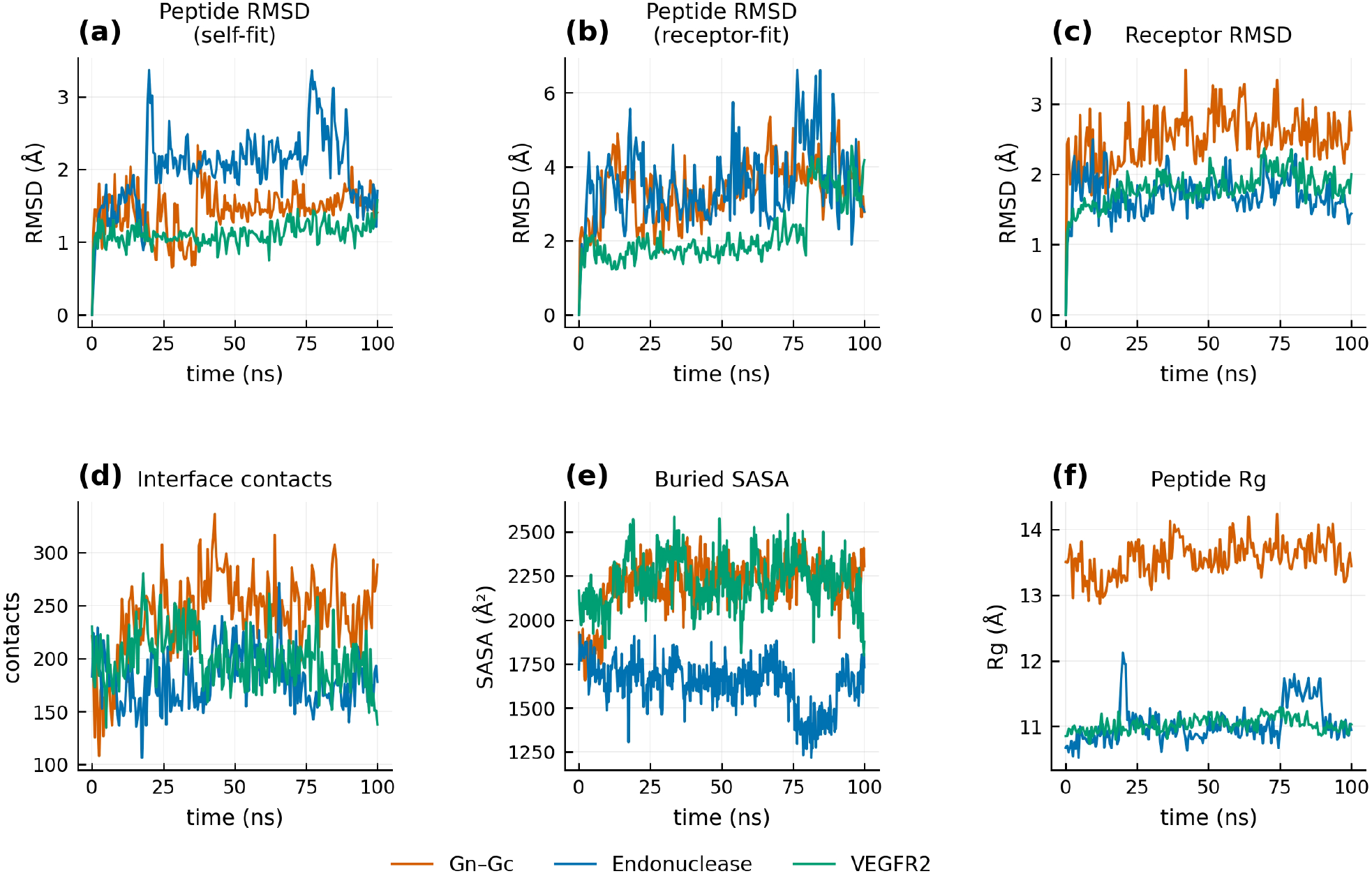
Molecular-dynamics stability over 100 ns for the three leads (Gn-Gc orange, 6Q99 blue, 2XIR green): (a) peptide Cα RMSD (self-fit), (b) peptide Cα RMSD (receptor-fit), (c) receptor Cα RMSD, (d) interface heavy-atom contacts, (e) buried SASA, (f) peptide radius of gyration.

**Figure 9.**
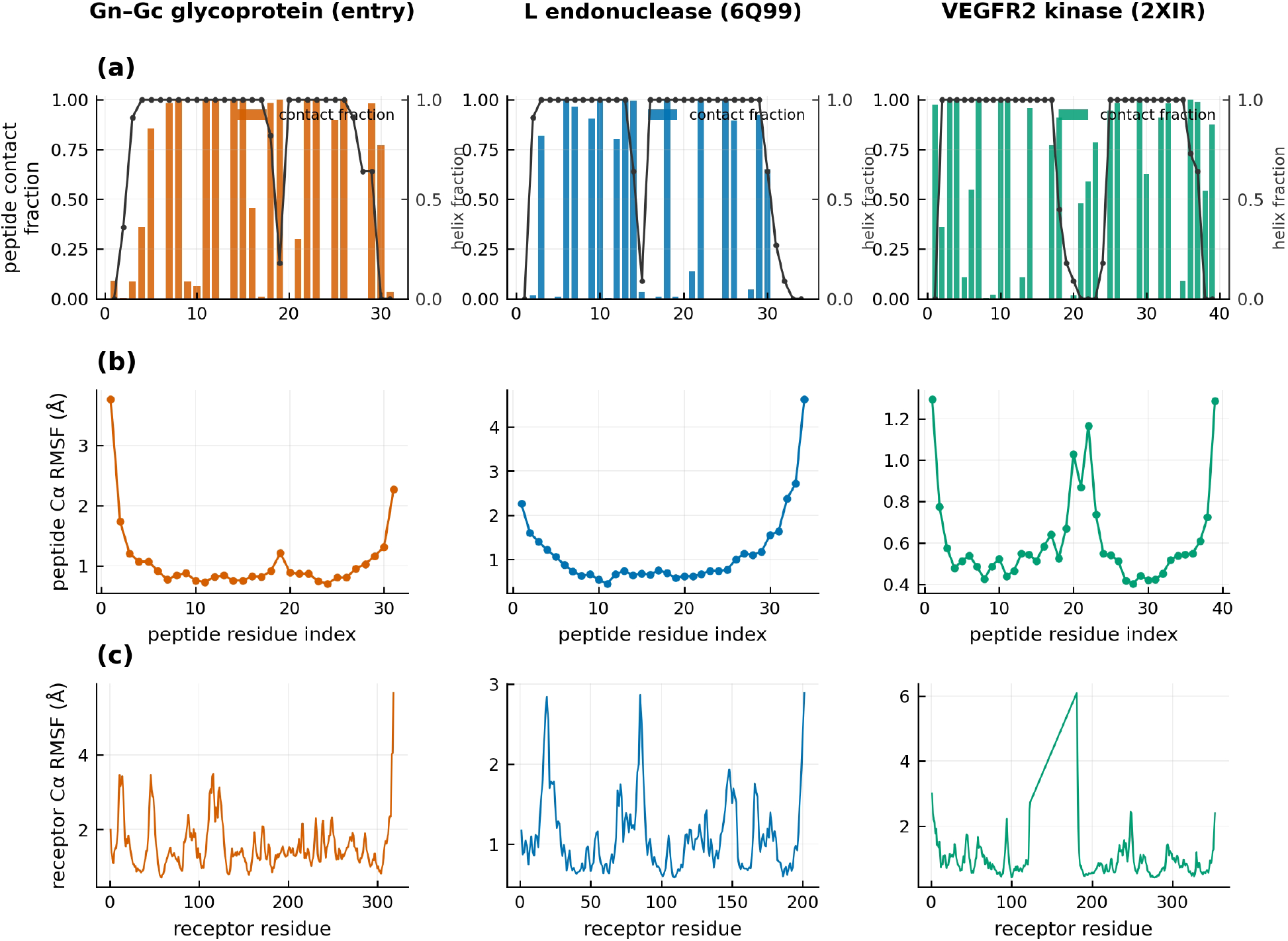
Per-residue interface characterization: (a) peptide interface contact occupancy (bars) with helix fraction (line), (b) peptide Cα RMSF (self-fit), (c) receptor Cα RMSF, for Gn-Gc, 6Q99, and 2XIR (columns). The 2XIR receptor RMSF peak near residues 124–180 reflects the modelled deletion junction.

The Gn-Gc binder not only retained but *expanded* its interface over the trajectory (contacts 221 → 288), consistent with productive relaxation into the binding pose. The 6Q99 binder was the most stable (interface essentially constant at 183.7 ± small fluctuations; receptor RMSD 1.72 Å). The 2XIR binder retained its fold tightly (self-fit RMSD 1.13 Å) and kept 100% contact retention even as its interface remodeled (230 → 138 contacts), reflecting redistribution rather than dissociation; its elevated final receptor-fit RMSD (4.18 Å at the last frame) localizes to the modelled 124–180 deletion junction rather than to the interface.

## 4. Discussion

### The platform view

Each stage of UnibindMD supplies an independent line of evidence, and the agreement across stages, rather than any single metric, is the basis for confidence. For the three ANDV leads: BindCraft predicted high-confidence helical interfaces (ipTM 0.87–0.94); PRODIGY independently predicted nanomolar-to-submicromolar ΔG (−9.1 to −11.1 kcal mol^−1^); ADMET-AI confirmed the expected peptide developability profile; and, decisively, 100 ns of explicit-solvent MD showed all three binders stably bound with retained helicity and ≥ 99.5% interface-contact retention. No single stage alone supports the claim; their consensus does.

### Target tractability

The endonuclease (6Q99) was the most designable (concave, well-defined catalytic site), whereas the flat kinase surface (2XIR) was least tractable yet yielded the highest-scoring individual designs, a pattern consistent with the broader de novo design literature [13,25] and with the observation that difficulty does not preclude quality when solutions are found.

### Design-strategy guidance

The non-monotonic hotspot–yield relationship (over-specification can reduce yield) suggests future campaigns should run systematic single-hotspot scans and then combine productive residues, rather than committing to many-constraint campaigns up front.

### Limitations

All results are computational and await experimental validation (cell-based entry-inhibition assays are the tractable first route; live-virus work requires BSL-3). The Boltz-2 cross-validation stage is incomplete owing to a missing ‘cuequivariance_torch’ dependency in the offline compute-node environment (MSA generation succeeded; inference did not). The 2XIR receptor carries a modelled deletion (124–180) whose junction dominates that system’s flexibility. ADMET-AI is a screening approximation, not a substitute for pharmacokinetic assays.

### Generalizability

Although demonstrated on ANDV, every stage of UnibindMD is target-agnostic: the pipeline accepts any protein (or protein complex) for which a structure exists or can be predicted, making it directly reusable for other antiviral, oncological, or antimicrobial binder programs.

## 5. Conclusion

We present **UnibindMD**, an integrated BindCraft–AlphaFold3-class–MD platform that takes a protein target from de novo peptide design through independent structure prediction, affinity and developability profiling, to physics-based dynamics validation. Applied to three ANDV intervention points, the platform generated 464 accepted binders and converged on three leads (ipTM 0.87–0.94; PRODIGY ΔG −9.1 to −11.1 kcal mol^−1^) that all remained stably bound over 100 ns of unrestrained molecular dynamics. By turning single-prediction design into a cross-validated, dynamics-tested workflow, UnibindMD provides a rapid, scalable, and generalizable computational path from viral (or any) target to validated peptide-binder candidates, directly addressing the urgent, currently unaddressed need for ANDV countermeasures.

## Data and Code Availability

All code, design trajectories, accepted sequences, analysis scripts, and MD trajectories supporting this work are available from the authors upon reasonable request, to ensure full reproducibility.

## Author Contributions

X.W. performed conceptualization, methodology, software development, and wrote the manuscript; A.H. contributed data curation and formal analysis; C.F. contributed investigation and validation; Z.W. contributed visualization; W.F. supervised the work, acquired funding, and is the corresponding author. All authors reviewed the manuscript.

## Acknowledgements

This work was supported by Shenzhen UnibindMD Computation Biotech Ltd, Co. Computational resources (including GPUs and the high-performance computing cluster) were provided by the UnibindMD supercomputing platform.

