## Supplementary figures for "UnibindMD: An Integrated AI and Molecular Dynamics Platform for De Novo Peptide Binder Design"

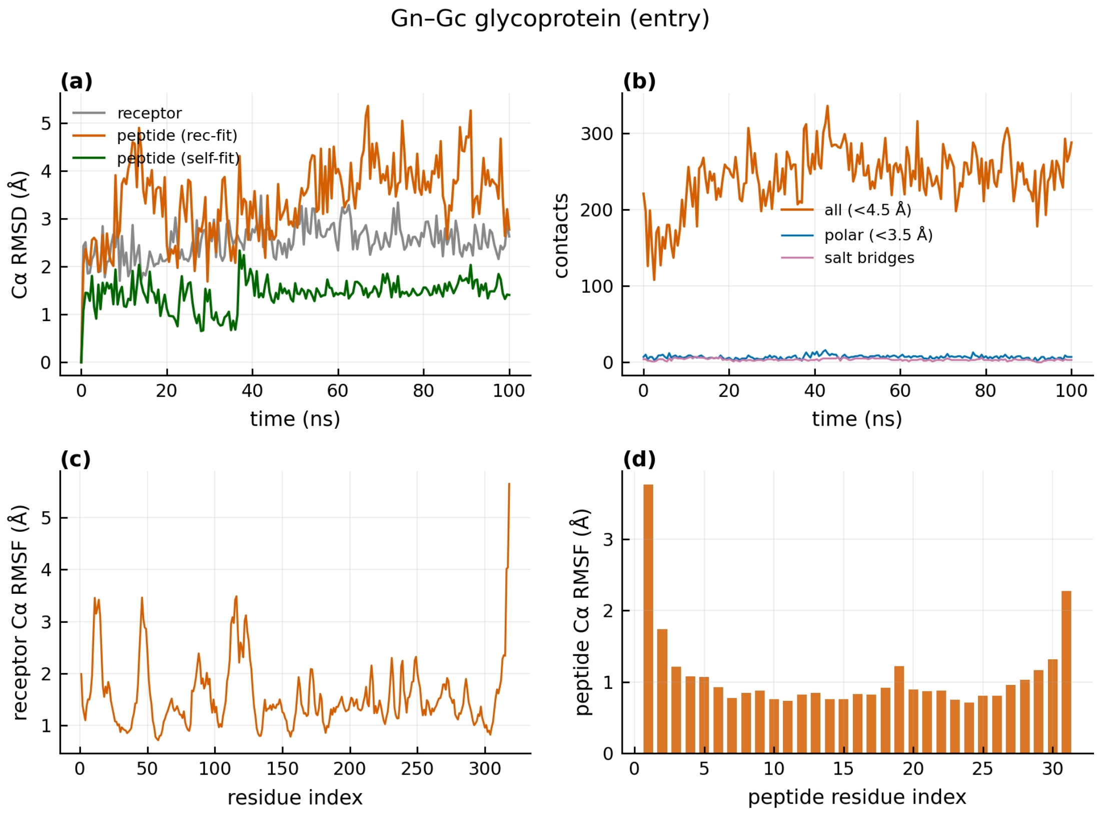


**Figure S1a.** Gn-Gc MD overview: Cα RMSD, interface contacts, receptor and peptide Cα RMSF.

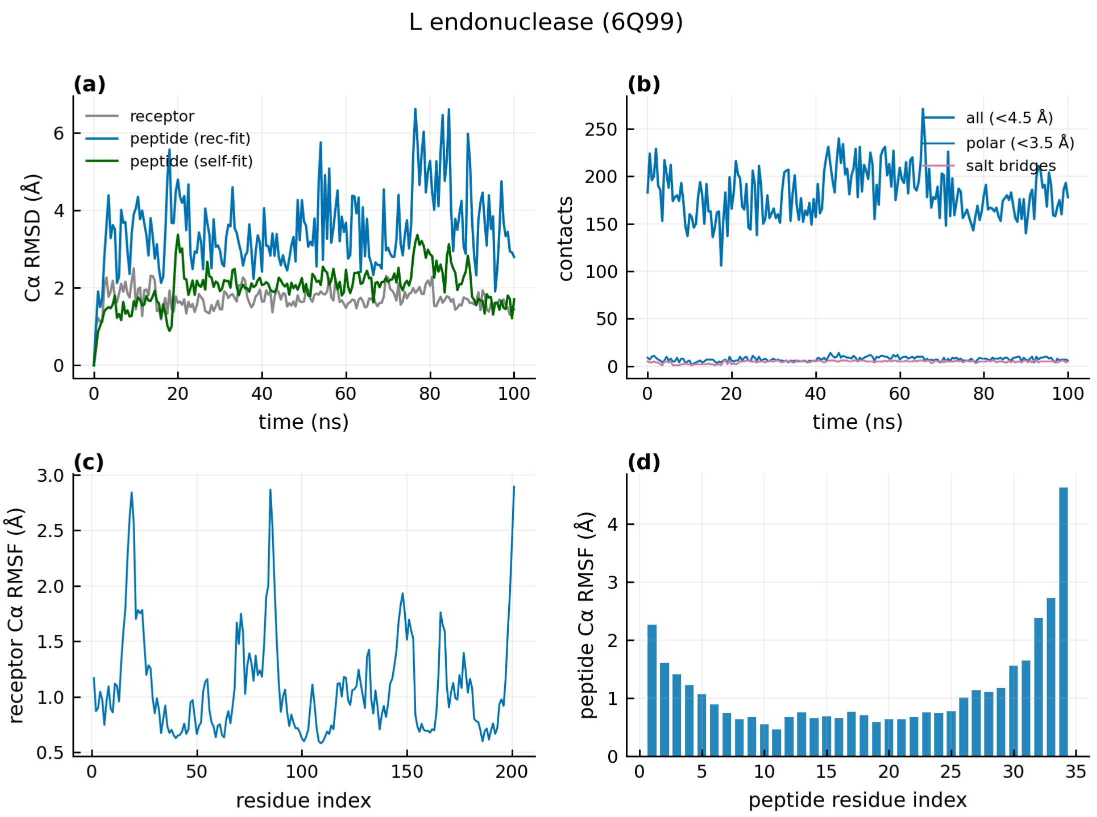


**Figure S1b.** 6Q99 MD overview: Cα RMSD, interface contacts, receptor and peptide Cα RMSF.

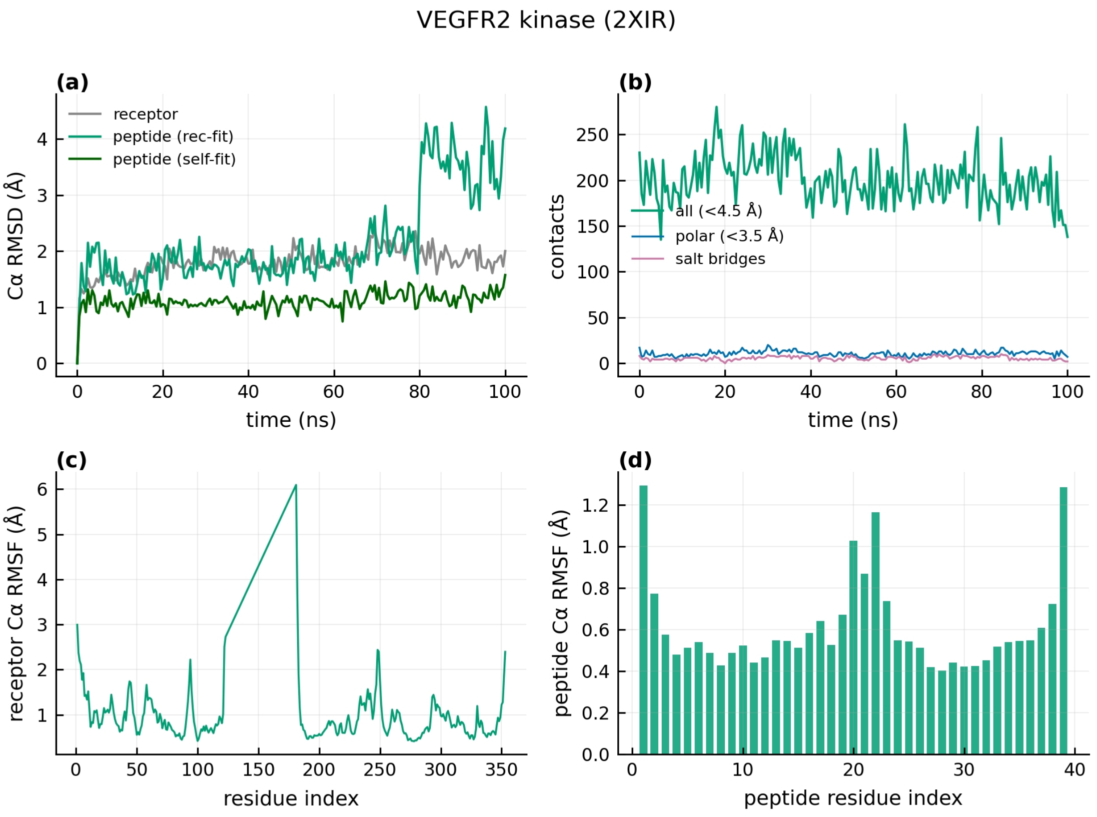


**Figure S1c.** 2XIR MD overview: Cα RMSD, interface contacts, receptor and peptide Cα RMSF.

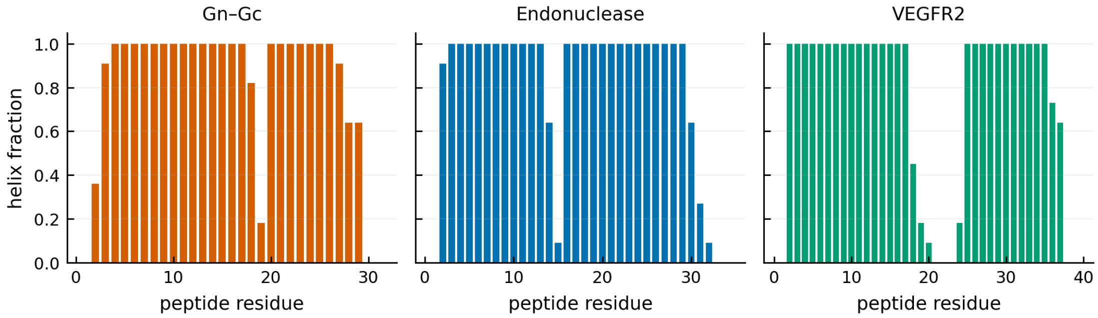


**Figure S2.** Per-residue helix fraction of the three peptide leads retained across the trajectory.

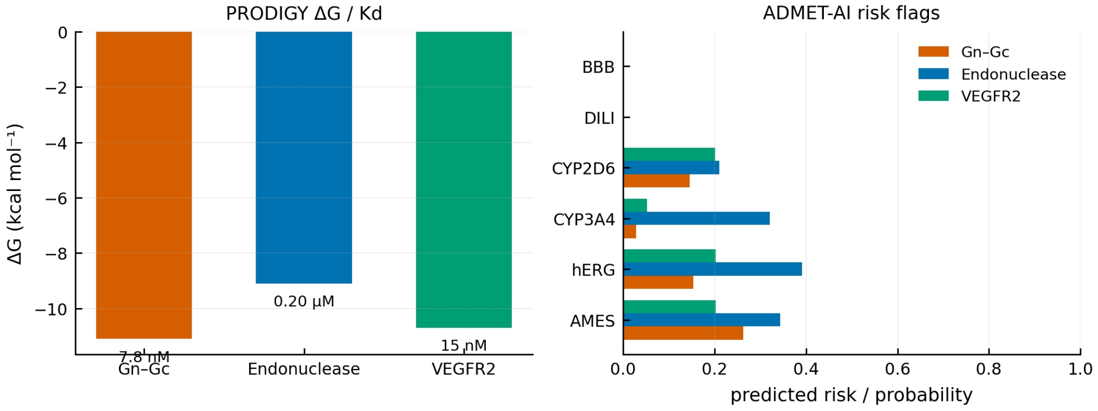


**Figure S3.** PRODIGY binding free energies (with Kd) and ADMET-AI risk-flag probabilities for the three leads.

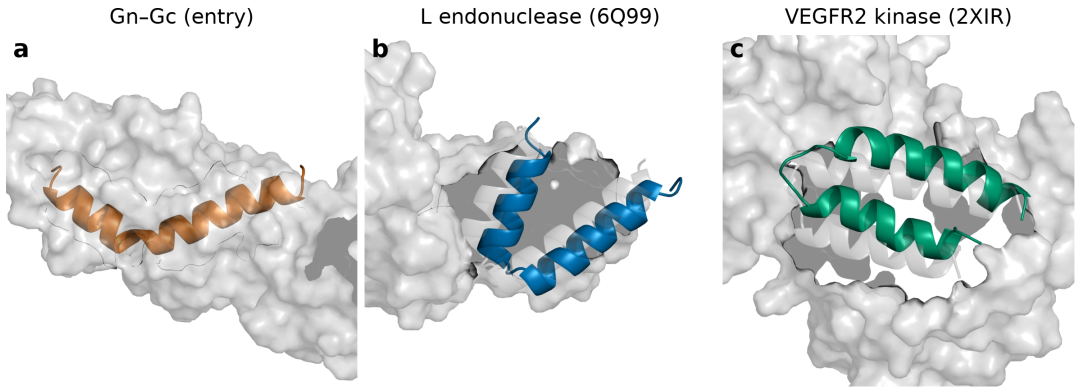


**Figure S4.** Design model (grey) vs MD final frame (coloured), aligned on receptor Cα: the binders retain their designed pose after 100 ns.
